# Microbial Associates of the Elm Leaf Beetle: Uncovering the Absence of Resident Bacteria and the Influence of Fungi on Insect Performance

**DOI:** 10.1101/2023.06.23.546294

**Authors:** Johanna Schott, Juliette Rakei, Mitja Remus-Emsermann, Paul Johnston, Susan Mbedi, Sarah Sparmann, Monika Hilker, Luis R. Paniagua Voirol

## Abstract

Microbial symbionts play crucial roles in the biology of many insects. While bacteria have been the primary focus of research on insect-microbe symbiosis, recent studies suggest that fungal symbionts may be just as important. The elm leaf beetle (ELB, *Xanthogaleruca luteola*) is a serious pest species of elm (*Ulmus minor*). Using culture-dependent and independent methods, we investigated the abundance and species richness of bacteria and fungi throughout various ELB life stages and generations, while concurrently analysing microbial communities on elm leaves. No persistent bacterial community was found to be associated with the ELB or elm leaves. By contrast, fungi were persistently present in the beetle’s feeding life stages and on elm leaves. Fungal community sequencing revealed a predominance of the genera *Penicillium* and *Aspergillus* in insects and on leaves. Culture-dependent surveys showed a high prevalence of two fungal colony morphotypes closely related to *Penicillium lanosocoeruleum* and *Aspergillus flavus*. Among these, the *Penicillium* morphotype was significantly more abundant on feeding-damaged compared to intact leaves, suggesting that the fungus thrives in the presence of the ELB. We assessed whether the detected prevalent fungal morphotypes influenced ELB’s performance by rearing insects on i) surface-sterilised leaves, ii) leaves inoculated with *Penicillium* spores, and iii) leaves inoculated with *Aspergillus* spores. Insects feeding on *Penicillium*-inoculated leaves gained more biomass and tended to lay larger egg clutches than those consuming surface-sterilised leaves or *Aspergillus*-inoculated leaves. Our results demonstrate that the ELB does not harbour resident bacteria and that it might benefit from associating with *Penicillium* fungi.

**Importance:** Our study provides insights into the still understudied role of microbial symbionts in the biology of the ELB, a major pest of elms. Contrary to expectations, we found no persistent bacterial symbionts associated with the ELB or elm leaves. Our research thus contributes to the growing body of knowledge that not all insects rely on bacterial symbionts. While no persistent bacterial symbionts were detectable in the ELB and elm leaf samples, our analyses revealed the persistent presence of fungi, particularly *Penicillium* and *Aspergillus* on both elm leaves and in the feeding ELB stages. Moreover, when ELB were fed with fungus-treated elm leaves, we detected a potentially beneficial effect of *Penicillium* on the ELB’s development and fecundity. Our results highlight the significance of fungal symbionts in the biology of this insect.

## Introduction

Microbial symbionts, here referred to as non-pathogenic microorganisms living in close association with a host, are harboured by many insect species. These symbiotic microorganisms influence diverse aspects of insect biology, such as nutrition, development, reproduction, immunity, and responses to abiotic stress (Engel and Moran 2013; Douglas 2015; Lemoine *et al*. 2020). The close associations of microbes with insects are subjected to complex co-evolutionary processes requiring fine-tuned adaptations from both the host and the symbiont (Kaltenpoth *et al*. 2014; Moran *et al*. 2019).

Although numerous insects have been demonstrated to depend on microbial symbionts, recent studies have questioned the prevailing assumption that all insects rely on microbial symbionts (Hammer *et al*. 2019). For example, stick insects (Phasmatodea) and caterpillars (Lepidoptera) do not harbour resident gut bacterial communities, suggesting that digestion and nutrition of these phyllophagous species do not depend on gut bacteria (Shelomi *et al*. 2013; Hammer *et al*. 2017; Paniagua Voirol *et al*. 2020b). As our understanding of the role of gut microbes in insects progresses, it has become clear that the extent to which insects depend on microbial symbionts varies widely within a broad range. This spectrum spans from a virtual absence of microbial symbionts to obligate mutualisms (Moran *et al*. 2019). However, since only a small fraction of insect species has been investigated for their microbial symbionts, it remains unclear how widespread associations with a resident microbial community are within the taxon Insecta.

To date, apart from studies on fungus-farming insects (Li *et al*. 2021) and termite gut protozoa (Brugerolle and Radek 2006), the majority of research on insect-microbe associations has focused on bacteria. Bacterial symbionts have been demonstrated to play a vital part in many hemimetabolous (Feng *et al*. 2019; Ohbayashi *et al*. 2019) and holometabolous insect species (Rio *et al*. 2012; Kwong and Moran 2016; Lavy *et al*. 2020; Moreau 2020; Hammer *et al*. 2023). However, our understanding of the impact of non-bacterial symbionts on insects is lacking behind. Only recently have interactions between phyllophagous insects and fungi started to receive more and more attention. For instance, a study by Berasategui *et al*. (2022) revealed a mutualistic relationship between the phytopathogenic fungus *Fusarium oxysporum* and the leaf beetle *Chelymorpha alternans*. In this interaction, the fungus protects the beetle’s pupal stage against predation, and in turn the beetle disperses the fungus to its host plant. Such findings suggest that fungal symbionts may be just as important as bacteria in influencing insect biology.

Beetles (Coleoptera) exhibit a broad spectrum of interactions with microbial symbionts (Salem and Kaltenpoth 2022). For instance, the burying beetle *Nicrophorus vespilloides* relies on gut bacteria to preserve its nutritional resources and enhance resistance against pathogens (Shukla *et al*. 2018; Wang and Rozen 2018). The tortoise leaf beetle *Cassida rubiginosa* depends on *Stammera* sp. in the gut; these bacteria provide digestive and detoxifying enzymes that facilitate nutrition from leaves (Salem *et al*. 2017; Salem *et al*. 2020). The bacterium *Burkholderia gladioli* protects the eggs of the darkling beetle *Lagria villosa* against fungal infection (Flórez and Kaltenpoth 2017; Niehs *et al*. 2020). Moreover, some symbionts can impact plant antiherbivore defences. An example for this trifold interaction, i.e. tritagonism (Freimoser *et al*. 2016), is the Colorado potato beetle *Leptinotarsa decemlineata*, harbouring oral bacteria that suppress plant antiherbivore defences (Chung *et al*. 2013).

Fungal symbiosis in beetles has predominantly been studied in bark beetles (Curculionidae). Bark beetles exhibit a diverse range of symbiotic relationships with fungi. Bark beetle-fungal interactions range from highly specialised fungus farming by beetles to weak interactions with fungal hitchhikers (Hulcr *et al*. 2020; Six 2020). As bark beetles feed on nutritionally poor substrates, their fungal partners often play a crucial role by providing important nutrients such as nitrogen and sterols (Klepzig and Six 2004). However, little is known on other types of beetle-fungus associations outside of the fungus-farming realm of beetles. Although beetles represent the most diverse insect taxon and encompass numerous pests that threaten crops and the forests alike, the interactions of most species with their microbial symbionts are understudied. Further research would advance our understanding of beetle ecology, enhance conservation strategies, and facilitate the development of pest management.

The elm leaf beetle *Xanthogaleruca luteola* (ELB; Chrysomelidae: Galerucinae) is a noteworthy pest species causing extensive damage to elms. As a specialised insect native to Europe and invasive in North America and Australia, it is a threat to elm populations (*Ulmus* spp.) (Dominiak and Kidston 2022). The voracious feeding on elm leaves by both, larval and adult stages, of the ELB results in widespread defoliation, stunted growth, increased susceptibility to pathogens, and ultimately tree mortality (Cranshaw and Zimmerman 2018). Despite the ecological and economic importance of the beetle’s biology, knowledge about its microbiota is lacking. Here, we investigated the abundance and species richness of bacteria and fungi across different life stages and generations of the ELB and its host plant (Figure 1). Moreover, we answered the question of whether they affect insect performance (Figure 2).

**Figure 1.**
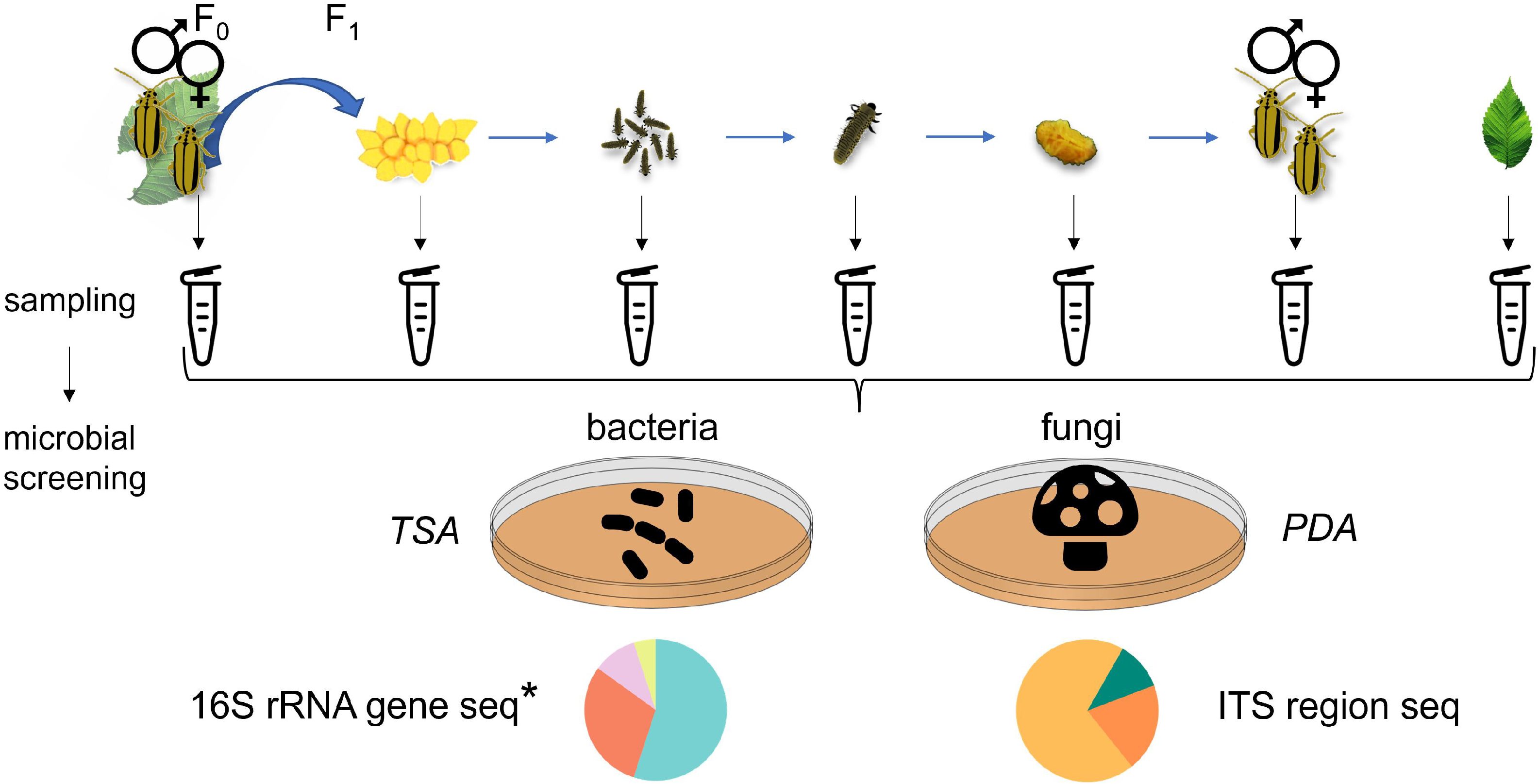
Overview of sampling and analysis of bacterial and fungal communities from different life stages and generations of the elm leaf beetle (*Xanthogaleruca luteola*) and from elm (*Ulmus minor*) leaves. Samples were collected from F_0_ beetles, their F_1_ eggs, neonate larvae, seven-day-old larvae, pupae, and freshly emerged F_1_ adults. We also sampled intact and beetle-fed elm leaves. Each sample was homogenised and divided into three parts: one part was plated on Trypticase Soy Agar (TSA), and another part was plated on Potato Dextrose Agar (PDA) for culture-dependent analysis of the bacterial and fungal communities. A third part was used for culture-independent analysis of fungal communities of the samples by ITS region sequencing. *For culture-independent analysis of the bacterial communities of the samples by 16S rRNA gene sequencing, an independent experiment following the same design was conducted.

**Figure 2.**
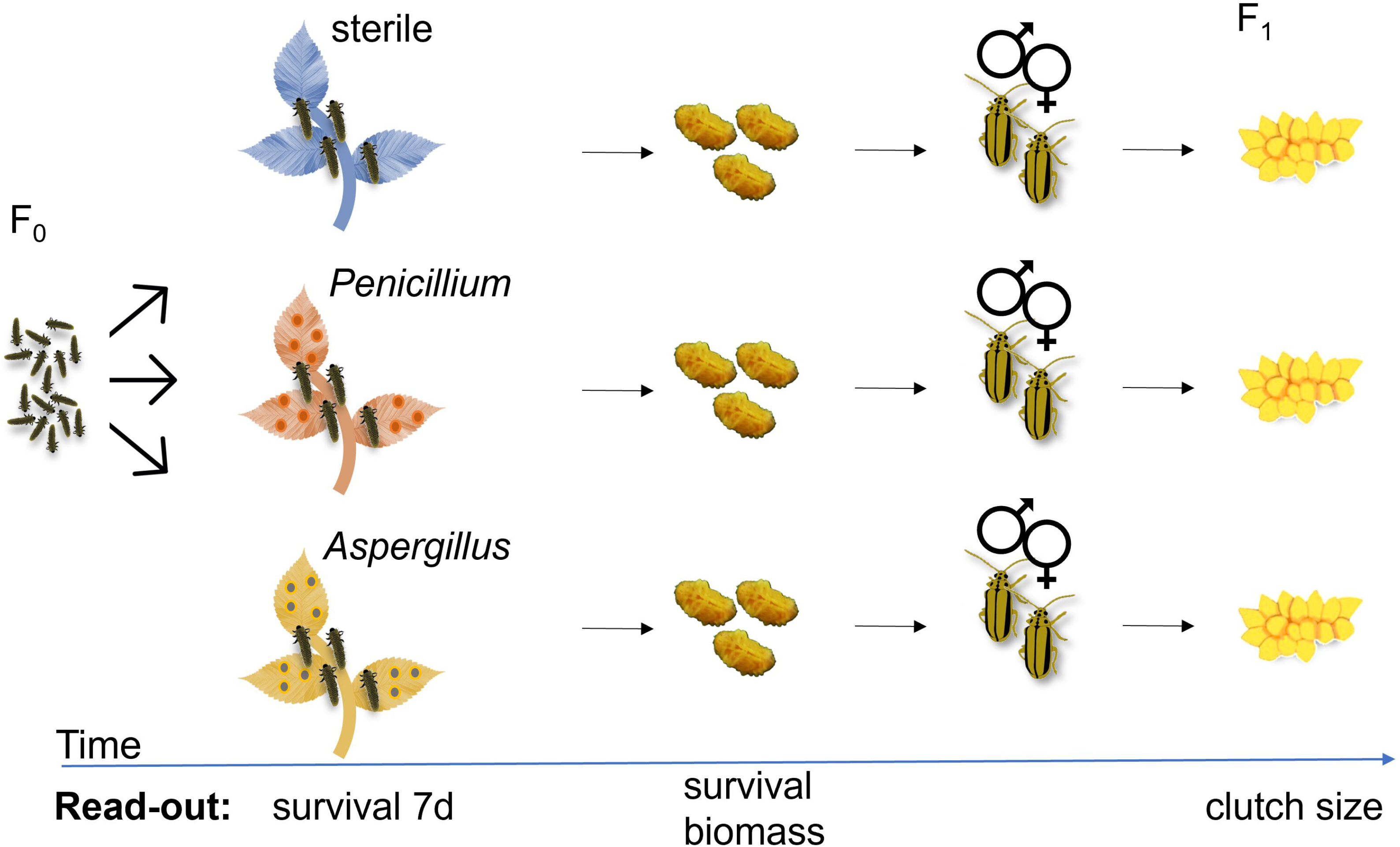
Assessing the effect of fungi on the performance of the elm leaf beetle (*Xanthogaleruca luteola*). To examine the impact of *Penicillium* and *Aspergillus* on insect performance, neonate larvae were divided into three treatment groups: (i) feeding on surface-sterilised leaves, (ii) feeding on leaves inoculated with *Penicillium* spores, or (iii) feeding on leaves inoculated with *Aspergillus* spores. Each biological replicate comprised a group of five to eight neonates. The larval survival was recorded on day seven and until pupation. Pupal biomass was recorded. Upon emergence, adults from the same treatment were paired and placed on an untreated elm branch for mating and egg deposition. The number of eggs per egg clutch laid by females that spent their juvenile development on the differently treated leaves was recorded for a period of two weeks.

## Results

### Scarce bacterial presence observed across elm leaf beetle life stages and on host plant leaves

Using a culture-dependent approach, we investigated whether the ELB harbours a persistent bacterial community by examining CFU abundance and bacterial identity from parental adult insects and their offspring across all life stages. Concurrently, we analysed samples from intact elm leaves and leaves that had been fed on by adult beetles to determine if the host plant impacts on the bacterial communities found within the insects, and if feeding activity by the insects impacts on the leaf-associated bacterial community (Figure 1). We used laboratory-reared insects deriving from a natural population, and greenhouse-grown, young elm trees.

We did not detect any CFU in samples from neonates, seven-day-old larvae, pupae, and freshly emerged F_1_ adults. Samples from F_0_ adults, their eggs, and leaves (intact and beetle-fed) sporadically showed CFU. On average, F_0_ adult samples exhibited 5.9 x 10^3^ CFU per sample, primarily due to high CFU counts in only 4 out of 28 samples (Figure 3). Eggs had on average 0.6 CFU per sample. Intact leaves had on average 73 CFU per sample, while beetle-fed leaves had on average 10 CFU per sample. Thus, only a small portion of the samples provided CFU (≤ 25% of F_0_-adult, egg and leaf samples, Figure 3). From these samples, we identified only three different colony morphotypes. Sequencing of the 16S rRNA gene revealed that these morphotypes represented the genera *Pseudomonas*, *Serratia*, and *Acinetobacter* (see Table S1 in supplemental material). Overall, our culture-dependent analysis did not provide evidence of a persistent bacterial community in the ELB or on host plant leaves.

**Figure 3.**
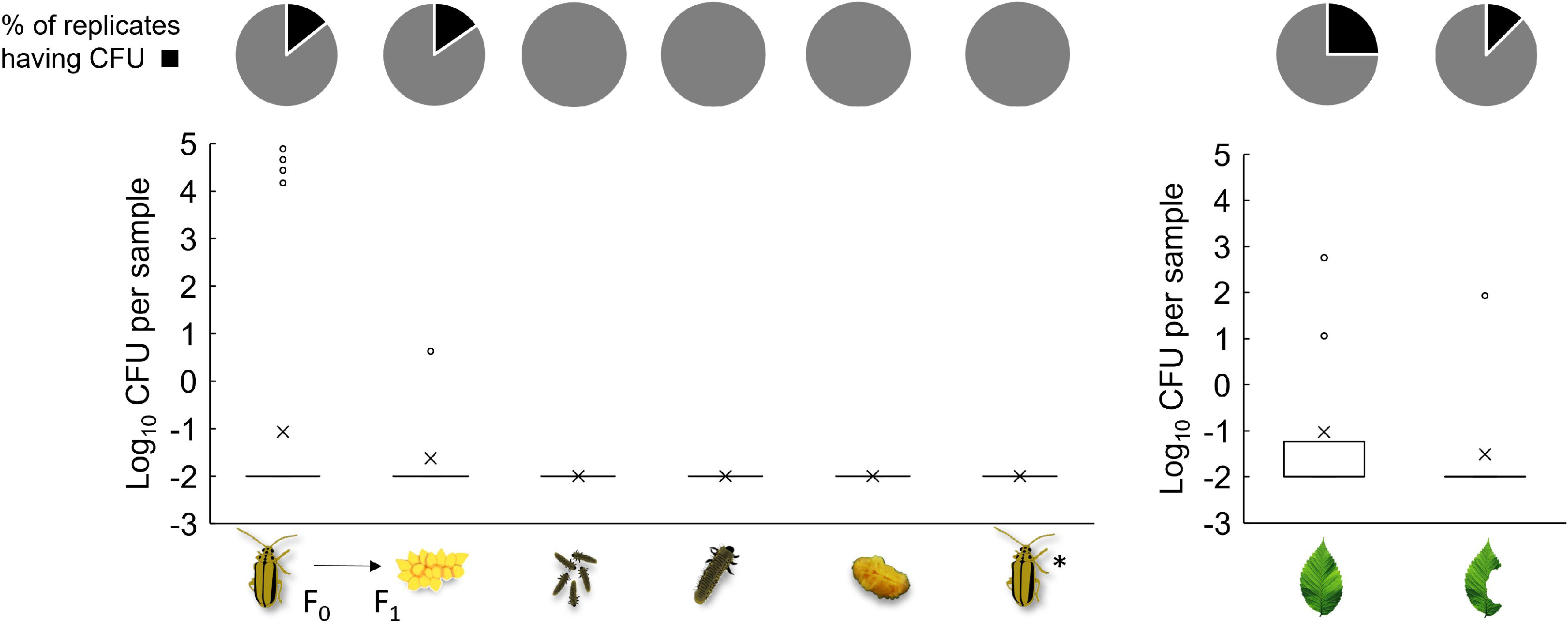
Culture-dependent analysis of bacterial abundance in elm leaf beetles (*Xanthogaleruca luteola*) and on host plant leaves (*Ulmus minor*). Pie charts depict presence (black) and absence (grey) of bacterial colony forming units (CFU) observed upon plating samples from F_0_ adults (n = 28), F_1_ eggs from different females (n = 14), pooled sibling neonates (n = 9), seven-day-old larvae (n = 8), pupae (n = 8), F_1_ adults (n = 16), intact leaves (n = 8), and beetle-fed leaves (n = 8) on Trypticase Soy Agar (TSA) medium. The boxplots display the distribution of log_10_(x + 0.01)-transformed CFU counts across beetle and leaf samples. The box in the boxplot represents the interquartile range (IQR), which contains the middle 50% of the data. The line inside the box indicates the median, while the × mark denotes the mean. Outliers are depicted as individual points. *F_1_ adults were collected upon emergence and had no contact with elm leaves.

To assess the extent to which the results obtained via culture-dependent methods are consistent with those obtained by a culture-independent approach, we conducted an independent experiment and performed 16S rRNA gene amplicon sequencing of samples obtained from adult insects, their offspring (all life stages) and intact elm leaves.

Our results show that most of the PCR products yielded bands that were either faint or undetectable upon electrophoresis (see Figure S1 in supplemental material). Nevertheless, the amplicons were sequenced. The sequencing results revealed that most bacterial reads detected in the insect and leaf samples were also present in the negative controls, suggesting that these bacterial amplicons originated from the so-called "kitome" (Salter *et al*. 2014; Paniagua Voirol *et al*. 2020a), and thus, were likely contaminants (Figure 4A). In accordance with the culture-dependent approach, the most abundant bacterial reads belonged to *Pseudomonas, Serratia,* and *Acinetobacter*. A principal coordinate analysis based on Bray-Curtis dissimilarity showed a separation between certain insect sample types (i.e., F_0_ adults, F_1_ eggs and adults) and negative controls or leaves. Nonetheless, this separation—observed in six out of 45 comparisons—was consistent with varying abundances of *Pseudomonas*, *Serratia*, and *Acinetobacter* reads, detected also in the negative controls, but not consistently present throughout the biological samples (Figure 4B). Thus, congruently with our culture-dependent analysis, the culture-independent analysis did not provide any evidence of a persistent bacterial community in the ELB or its host plant.

**Figure 4.**
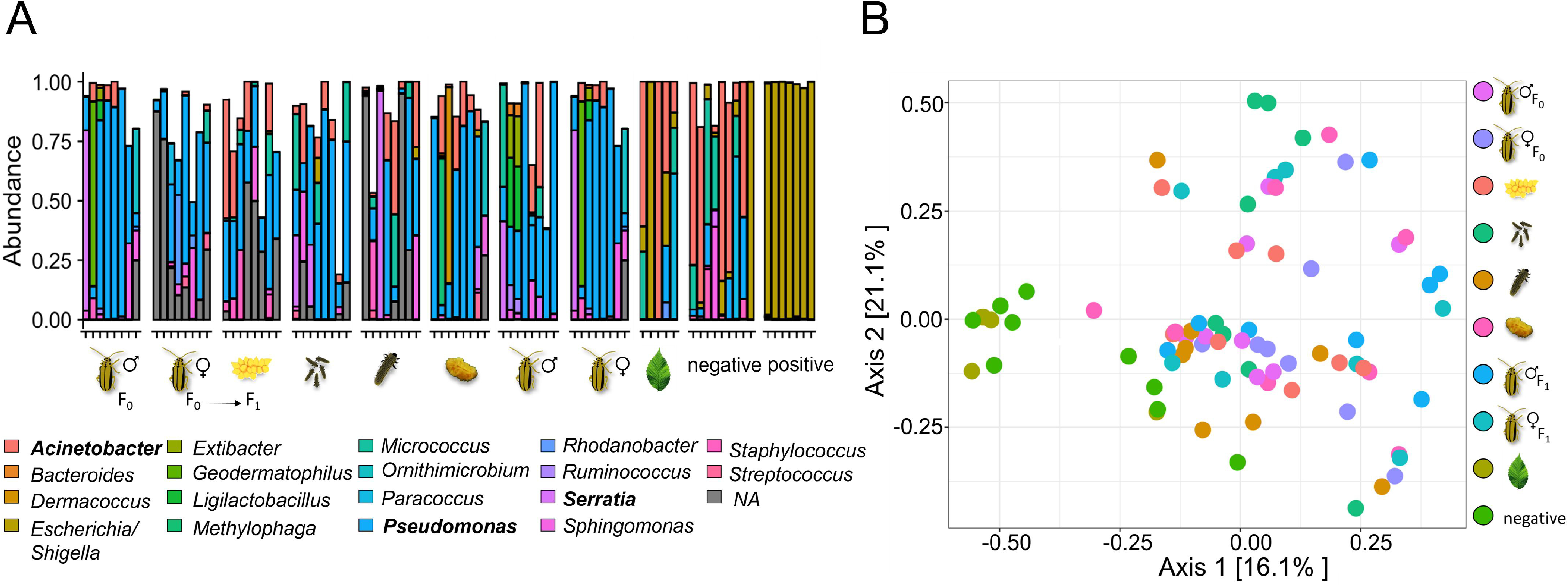
Culture-independent analysis of bacterial communities associated with different elm leaf beetle (*Xanthogaleruca luteola*) life stages and with host plant leaves (*Ulmus minor*). (A) Relative abundance of the 20 most abundant bacterial taxa, identified through MiSeq sequencing of the 16S rRNA gene. Two taxa lacked genus classification (NA). Each bar represents an individual sample. Three leaf samples of the originally 8 samples yielded fewer than 10 reads, and thus, were excluded. Genera identified through culture-dependent approaches are highlighted in bold. (B) Principal-coordinate (PCo) analysis of microbial beta diversity (based on Bray-Curtis dissimilarity) for insect samples, leaf samples, and negative controls. Pairwise comparisons of sample types showed significant differences between F_0_ female and F_1_ adult samples when compared to the negative controls. Similarly, egg, F_0_ male, and F_1_ male samples significantly differed from leaf samples (P < 0.05, PERMANOVA).

### High abundance of fungi in feeding insect life stages and beetle-fed leaves

To investigate the abundance and species richness of fungi in ELBs and on elm leaves, we also followed a culture-dependent and independent approach. CFU and fungal identity from samples of parental adult insects and their offspring across all life stages, as well as of intact leaves and beetle-fed leaves (Figure 1) were determined. We found that samples from non-feeding insect stages showed only sporadic presence of fungal CFU (≤ 25% of the samples, Figure 5). By contrast, most insect samples from feeding stages and both intact and beetle-fed leaves yielded CFU (≥ 96% of the samples, Figure 5). Notably, no CFU were obtained from samples of ELB pupae. Samples from F_0_ adults and F_1_ larvae yielded significantly more CFU than F_1_ eggs, F_1_ neonates, F_1_ pupae and freshly emerged F_1_ adults, which did not feed yet (Figure 5). Moreover, we observed that beetle-fed leaves carried on average 30 times more CFU than intact leaves (Figure 5). Hence, our analysis revealed conspicuous fungal presence during the feeding life stages of the beetles and in feeding-damaged elm leaves.

**Figure 5.**
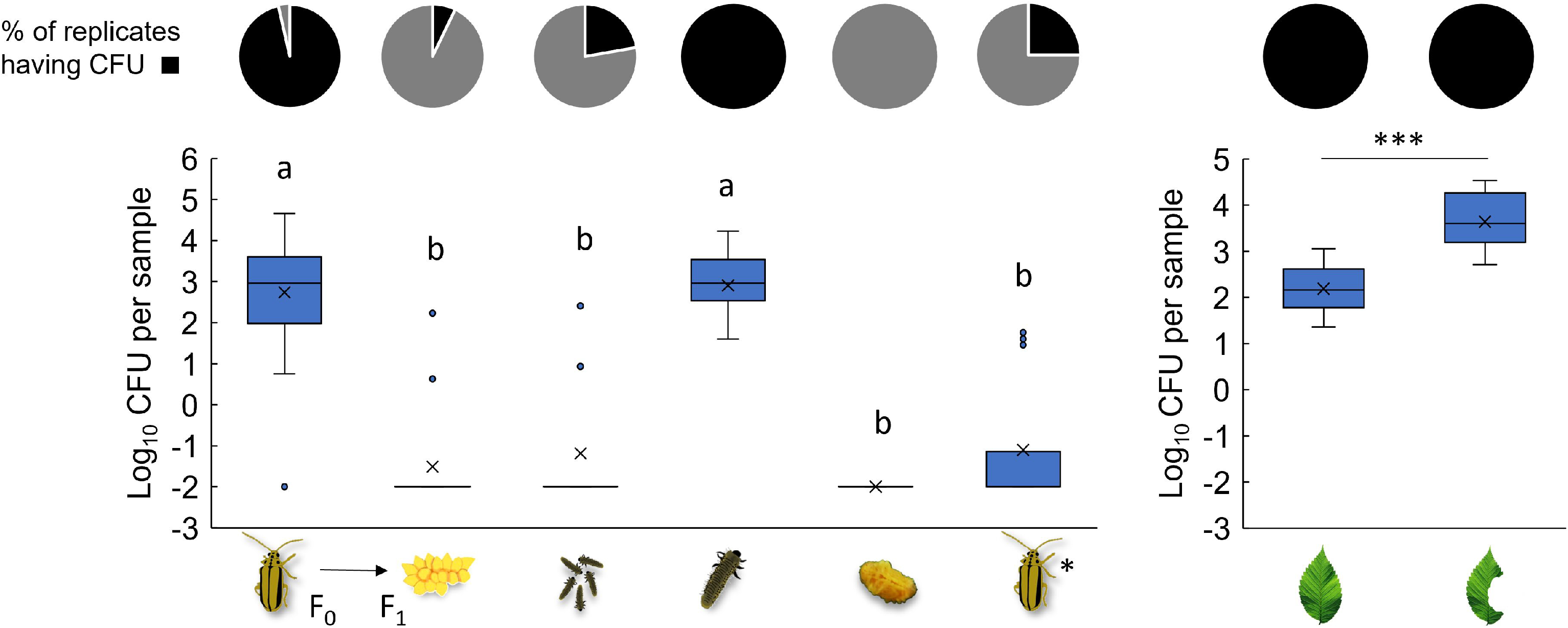
Culture-dependent analysis of fungal abundance in elm leaf beetles (*Xanthogaleruca luteola*) and on host plant leaves (*Ulmus minor*). Pie charts depict presence (black) and absence (grey) of fungal colony forming units (CFU) observed upon plating samples from F_0_ adults (n = 28), F_1_ eggs from different females (n = 14), pooled sibling neonates (n = 9), larvae (n = 8), pupae (n = 8), F_1_ adults (n = 16), intact leaves (n = 8), and beetle-fed leaves (n = 8) on Potato Dextrose Agar (PDA) medium. The boxplots display the distribution of log_10_(x+0.01)-transformed CFU counts across beetle and leaf samples. Boxes represent the interquartile range (IQR) with the median (line) and mean (×) inside. Whiskers extend to data points within 1.5 times the IQR, and outliers are shown as individual points. Different letters or asterisks above the bars indicate significant differences between groups (insects: KW test p < 0.001; Dunn-BH p < 0.001; leaves: *t*-test p < 0.001). *F_1_ adults were collected upon emergence and had no contact with elm leaves.

We identified two different colony morphotypes. Sequencing of the internal transcribed spacer (ITS) region of the ribosomal operon revealed that these morphotypes were closely related to *Penicillium lanosocoeruleum* and *Aspergillus flavus* (see Table S2 in supplemental material). These morphotypes were designated as *Penicillium* sp. LPV01 and *Aspergillus* sp. LPV02, respectively. Their colonies were visually distinguishable, enabling us to enumerate their respective abundances in samples from F_0_ adults and F_1_ larvae, as well as on intact and beetle-fed leaves. In samples from F_0_ adult beetles, *Penicillium* was significantly more abundant than *Aspergillus*. *Penicillium* abundance in F_0_ adults was comparable to that of F_1_ larvae. Likewise, *Aspergillus* abundance in samples from F_0_ adult beetles was comparable to that in samples from F_1_ larvae (Figure 6).

**Figure 6.**
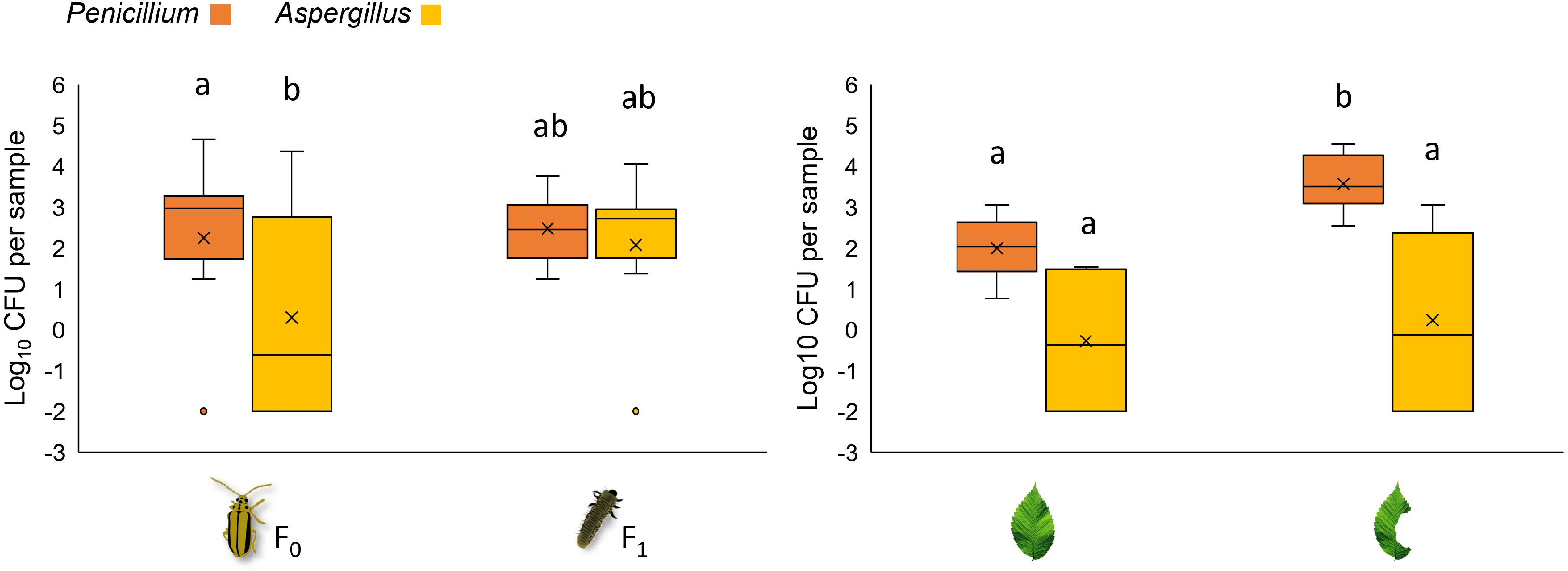
Colony forming units of *Aspergillus* sp. LPV01 and *Penicillium* sp. LPV02 obtained from samples of the feeding stages of the elm leaf beetle (*Xanthogaleruca luteola*) as well as of intact elm (*Ulmus minor*) leaves, and beetle-fed elm leaves. The boxplots display log-transformed colony forming unit (CFU) counts in samples from the elm leaf beetle feeding stages (F_0_ adults, n = 28; F_1_ larvae, n = 8), intact leaves (n = 8), and beetle-fed leaves (n=8) on Trypticase Soy Agar (TSA) medium. Boxes represent the interquartile range (IQR) with the median (line) and mean (X) inside. Whiskers extend to data points within 1.5 times the IQR, and outliers are shown as individual points. Different letters above bars indicate significant differences (insects: KW test p = 0.002; Dunn-BH p < 0.01; leaves: KW test p < 0.001 Dunn-BH: p < 0.05).

When comparing CFU from intact and beetle-fed leaves, we observed marginally significant differences between *Aspergillus* and *Penicillium* counts in intact leaves, with *Penicillium* sp. LPV01 exhibiting higher numbers (p = 0.054). *Aspergillus* was similarly abundant in beetle-fed leaves and intact leaves, whereas the *Penicillium* population increased by 35-fold on beetle-fed leaves (Figure 6). These findings suggest that the rise in total CFU counts in beetle-fed leaves is primarily attributed to the increase in the *Penicillium* population rather than *Aspergillus*. Hence, *Penicillium* appears to play a significant role in shaping the fungal community fungal community within the beetle - elm interaction, with its proliferation being particularly pronounced in beetle-fed leaves.

To investigate the potential occurrence of fungal taxa that might not be culturable, we performed an amplicon sequencing of the ITS of the ribosomal operon. Our findings revealed that the *Penicillium* genus was the most abundant one in insect samples (F_0_ parents and their offspring across all life stages). *Aspergillus* was the second most abundant genus, identified across all insect life stages, albeit not in every sample. By contrast, the most abundant reads in both intact and beetle-fed leaf samples were attributed to *Aspergillus*. Notably, *Penicillium* was detected on beetle-fed leaves (Figure 7). Thus, in accordance with our culture-dependent analysis, our culture-independent screening indicates that the fungal community associated with the ELB, and elm leaves is dominated by *Penicillium* and *Aspergillus*.

**Figure 7.**
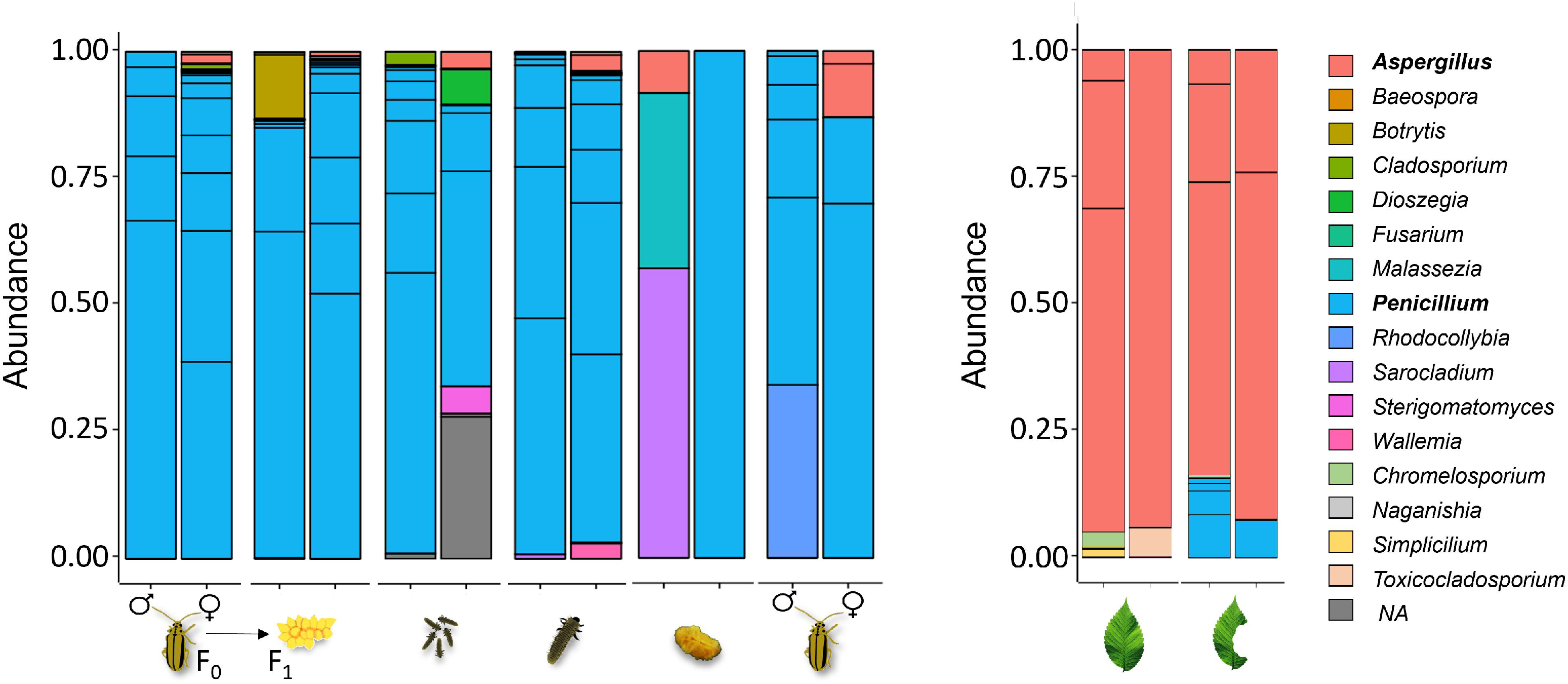
Culture-independent analysis of fungal communities associated with different elm leaf beetle (*Xanthogaleruca luteola*) life stages and with host plant (*Ulmus minor*) leaves. Relative abundance of fungal taxa in beetles (left) and elm leaves (right), identified through MiSeq sequencing of the ITS rRNA gene. Three taxa lacked genus classification (NA), resulting in 17 displayed keys.

### Effect of *Penicillium* sp. LPV01 and *Aspergillus* sp. LPV02 on insect performance

We investigated whether the predominant fungi observed in the ELB active feeding stages and on elm leaves influence beetle performance. We reared the insects on three types of leaves: surface-sterilised leaves, leaves inoculated with *Penicillium* sp. LPV01 spores, and leaves inoculated with *Aspergillus* sp. LPV02 spores. Subsequently, we analysed larval survival at day seven, survival until pupation, and pupal biomass. After allowing the insects to reach adulthood and mate, we compared the egg clutch sizes produced by females that spent their juvenile development on the differently treated leaves; the females were offered untreated trees for depositing their eggs (Figure 2).

We found no significant differences in the survival rates of seven-day-old larvae fed on surface-sterilised leaves or on leaves inoculated with either type of fungus (Figure 8A). Similarly, pupal survival rates were not significantly affected by the different treatments (Figure 8B). On the other hand, the biomass of pupae significantly differed depending on the treatment (Figure 8C). Insects that had fed on *Penicillium*-inoculated leaves gained more biomass than those fed on surface-sterilised leaves and those fed on *Aspergillus*-inoculated leaves. In contrast, pupae that had fed on *Aspergillus*-treated and surface-sterilised leaves during their larval development had similar biomass (Figure 8C). We also observed marginally significant differences in sizes of egg clutches laid by females subjected to different treatments during their juvenile development (Figure 8D). Females that had developed on *Penicillium*-inoculated leaves tended to produce egg clutches with more eggs than those that had developed on sterile leaves or *Aspergillus*-inoculated leaves. The egg clutch sizes of females that had developed on sterile leaves and those on *Aspergillus*-inoculated leaves were not significantly different.

**Figure 8.**
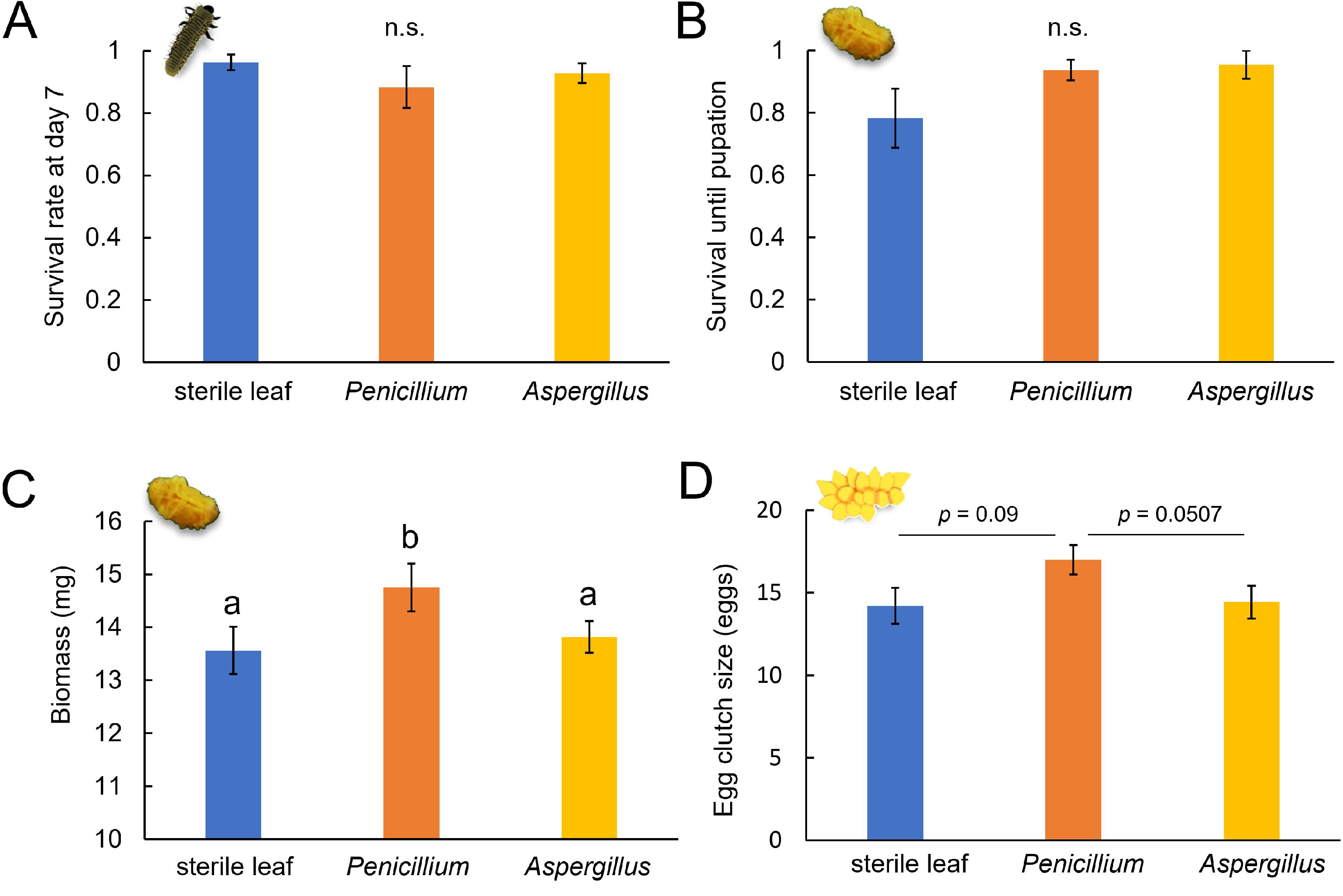
Effects of *Penicillium* and *Aspergillus* fungi on elm leaf beetle (*Xanthogaleruca luteola*) performance. Insects were reared on three types of elm (*Ulmus minor*) leaves: surface-sterilised leaves (blue), leaves inoculated with *Penicillium* spores (orange), and leaves inoculated with *Aspergillus* spores (yellow). We recorded (A) larval survival at day seven (n = 12), (B) pupal survival (n = 11-12), (C) pupal biomass (n = 11-12), and (D) egg clutch size (number of eggs per clutch) produced by females subjected to the different treatments (n = 15-17). Error bars represent standard error of the mean (SEM). Different letters above the bars indicate significant differences between groups (KW test; p < 0.05; Dunn-BH: p < 0.05), “n.s.” indicates no significant difference.

Overall, our findings suggest that *Penicillium* positively affects the ELB performance. By contrast, *Aspergillus* has no discernible effect on the ELB performance.

## Discussion

Our study showed that larvae and adults of the ELB take up fungi from host plant leaves during feeding, but the ELB does not vertically transfer these microbes. Ingested *Penicillium* spores from elm leaves exert some beneficial effects on the beetle’s performance. In contrast, no evidence of a persistent bacterial community was detected in the different life stages of the ELB or on the leaves of its host plant.

The scarcity of bacteria in the studied ELB life stages contrasts the presence of beneficial gut bacteria in various coleopteran species. For example, several Scarabaeidae, Cerambycidae, and Curculionidae are known to harbour bacteria with cellulase and xylanase activity (Banerjee *et al*. 2022). Similarly, the Colorado potato beetle (*Leptinotarsa decemlineata*) of the Chrysomelidae family was found to host gut bacteria with cellulase and xylanase activities. Furthermore, this species harbours bacteria capable of suppressing plant defences upon oral secretion into plant wounds (Vilanova *et al*. 2012; Chung *et al*. 2013). Additionally, gut bacteria of reed beetles (Chrysomelidae) contribute to the synthesis of essential amino acids and the production of the B vitamin riboflavin. They also support the digestion of host plant leaves through the production of pectinases (Reis *et al*. 2020).

The low presence of transient and lack of persistent environmental bacteria in the gut of the ELB and on elm leaves is surprising, especially when considering that the insect gut typically offers favourable conditions for bacteria (Engel and Moran 2013; Moran *et al*. 2019). Likewise, the phyllosphere is well known to be colonised by cultivatable bacteria (Remus-Emsermann and Schlechter 2018). A study on the gut microbiome of eleven species of *Longitarsus* flea beetles revealed that gut bacteria are likely acquired from the environment. The diversity of bacteria associated with *Longitarsus* had no correlation with the beetles’ phylogeny or life-history traits (Kelley and Dobler 2011). One possible reason for the low detection of bacteria in the ELB is the high content of flavonoids, such as kaempferol and quercetin, present in elm leaves (Schott *et al*. 2022). While many gut-associated bacteria in insects can break down flavonoids and other phenolic compounds (Cheng *et al*. 2018), it is worth noting that kaempferol and quercetin also have antimicrobial properties (Górniak *et al*. 2019). Moreover, it is known that egg deposition by the ELB can enhance the levels of kaempferol and quercetin in elm leaves (Schott *et al*. 2022). These antimicrobial compounds might impair bacterial growth in the ELB gut when plant cells are disrupted, and bacteria will be exposed to the flavonoids. Furthermore, the fungi present on the elm leaf surface might contribute to bacterial growth inhibition on elm leaves and, when leaves are ingested by the ELB, also in the ELB gut. *Penicillium* and *Aspergillus* species are well-known to produce antibacterial compounds (Fleming 1929; Frisvad *et al*. 2004; Al-Fakih and Almaqtri 2019; Khattak *et al*. 2021). Thus, it is tempting to speculate that the presence of these fungi suppresses bacterial growth in the ELB - elm system. Further research investigating the potential antibacterial effects of the retrieved strains could offer valuable insights into whether they inhibit bacterial growth on elm leaves and in the gut of the ELB.

In addition to the phytochemistry of elm leaves and the presence of fungi on the leaves, the semi-natural conditions of our experiments may have contributed to the limited presence of environmental bacteria. The phyllosphere microbiomes of plants grown under greenhouse conditions are known to differ from those grown in natural settings, with the former exhibiting lower microbial abundance and diversity (Williams and Marco 2014; Wei *et al*. 2016). However, plants cultivated under greenhouse conditions are usually also colonised by a number of bacteria in notable amounts (Maignien *et al*. 2014). Similarly, laboratory-reared insects were found to be colonised by fewer microbes and fewer microbial species than their wild counterparts (Belda *et al*. 2011; Staudacher *et al*. 2016). The beetles used in our study derived from a natural population. Essential gut or intracellular bacterial symbionts are expected to be retained and passed down through generations even when the ELB host is fed with greenhouse-grown leaf material. However, since we did not observe this, our study provides evidence that the successful development and reproduction of the ELB does not depend on the presence of bacterial associates.

The independence of the ELB performance of persistently present bacterial associates indicates that endogenous, self-produced enzymes are available for efficient digestion of elm leaves. Indeed, many herbivorous beetles are able to digest their host plants independently from microbial symbionts. A phylogenomic analyses showed that the Phytophaga clade, which includes the Chrysomeloidea superfamily, acquired plant cell wall degrading enzymatic activity through horizontal gene transfer (HGT) from microbes. This genetic adaptation enabled the beetles to exploit woody tissues and pectin-rich leaves independently of microbial symbionts (McKenna *et al*. 2019). Interestingly, a study by Kirsch *et al*. (2014) provided evidence suggesting that a pectin-degrading polygalacturonase encoding gene from an ascomycete fungus has been acquired by HGT by a common ancestor of Chrysomeloidea and Curculionidea. Gene duplications and further HGTs led to functional diversification of these digestive enzymes.

The ascomycete fungi *Penicillium* and *Aspergillus* fungi were found to be abundant in the ELB feeding stages and on elm leaves, while their presence was minimal in eggs, neonate larvae, pupae, and freshly emerged adults. These findings demonstrate that fungi are not transferred between different life stages or generations in the ELB. Instead, it is likely that the ELB acquires these fungi from the environment while feeding. Interestingly, when comparing beetle-fed leaves to intact leaves, we discovered a significant increase in fungal abundance. This increase was primarily attributed to a 35-fold higher abundance of *Penicillium* sp. LPV01 in beetle-fed leaves compared to intact leaves, suggesting that this fungus thrives in the presence of the ELB. One possible explanation for this phenomenon is that *Penicillium* sp. LPV01 obtains nutrients from the ELB’s faeces since fungal growth on faeces was frequently detected (personal observations). Alternatively, the fungus may utilise plant nutrients released when the beetle damages the leaves. Moreover, *Penicillium* sp. LPV01 could also multiply within the beetle’s gut upon ingestion and subsequently be excreted onto the leaves during defecation, contributing to the observed higher CFU numbers on beetle-fed leaves.

*Penicillium* sp. LPV01 exhibited the highest similarity to *P. lanosocoeruleum*, a fungus isolated from various plants and soil (Pitt *et al*. 1997; Samson *et al*. 2004; Debbarma *et al*. 2021; Zheng *et al*. 2022). Therefore, it is likely that *Penicillium* sp. LPV01 commonly colonises plant surfaces. However, it remains unclear what resources this fungus utilises on elm leaves in the absence of the ELB. Intact elm leaves colonised by *Penicillium* sp LPV01 were asymptomatic, indicating that this fungus is no phytopathogen of elm. *Aspergillus* sp. LPV02 exhibited the highest similarity to *A. flavus*, a fungus with saprophytic and pathogenic characteristics and widely found in soil, water, air samples, and both healthy and diseased plant tissue (Ramírez-Camejo *et al*. 2012). We found no indication that *Aspergillus* sp. LPV02 exerts phytopathogenic activity on elm leaves. The number of *Aspergillus* sp. LPV02 CFU did not increase after beetle damage. Therefore, unlike *Penicillium* sp. LPV01, there is no evidence suggesting that *Aspergillus* sp. LPV02 thrives in the presence of the ELB.

Our study on the impact of the detected *Penicillium* and *Aspergillus* morphotypes on the performance of the ELB suggests that the ELB benefits from taking up *Penicillium* sp. LPV01 with respect to pupal biomass gain in the end of the juvenile development. Moreover, the females resulting from the heavy pupae that developed on *Penicillium* treated leaves tended to lay more eggs per egg clutch than females that developed on untreated leaves. Consistent with our observations, a recent study showed a positive correlation between pupal mass and egg numbers in the early egg laying phase of the ELB (Schott *et al*. 2023).

However, it remains unclear how *Penicillium* sp. LPV01 contributes to the improved performance of ELB. The fungus might support the digestion of elm leaves by improving the degradation of plant cell wall components in the ELB gut. For example, *P. crustosum* and *Fusarium culmorum*, residing in the gut of the linden borer (*Saperda vestita*, Coleoptera: Cerambycidae), can degrade cellulose (Delalibera *et al*. 2005). Similarly, *F. solani* found in the gut of Asian longhorned beetles (*Anoplophora glabripennis*, Coleoptera: Cerambycidae), contributes to lignocellulose digestion (Geib *et al*. 2008; Wang *et al*. 2022). Moreover, *Penicillium* might improve the ELB performance by circumventing or suppressing the elm antiherbivore defences. Elms are known to increase the levels of the kaempferol and quercetin derivatives in their leaf tissue as a response to ELB infestation; the induced high concentration of a kaempferol derivative was shown to result in increased larval mortality (Austel *et al*. 2016; Schott *et al*. 2022). *Penicillium* sp. LPV01 might assist in the degradation of these defence compounds. Many *Penicillium* (and *Aspergillus*) strains are known to transform and metabolise flavonoids (Cao *et al*. 2015). Thus, it is reasonable to hypothesise that *Penicillium* sp. LPV01 influences the beetle’s susceptibility to flavonoid exposure.

### Conclusion

Our study adds to the growing body of research showing that fungal symbionts of phyllophagous beetles may play important roles in shaping interactions between these beetles and their host plants. Further research is needed addressing the metabolic abilities of the fungi, thus elucidating how they might support the beetles in leaf digestion. Such studies should not only focus on plant cell wall degradation activities of the fungus, but also take into account how the fungal symbiont changes defensive plant metabolites in the beetle’s gut, thereby probably mitigating plant defences. Moreover, future studies on the impact of fungal symbionts on the performance of the host should also consider that functions of different fungal species might interfere and shape the outcome. Additionally, genome analyses of the beetles and their transient fungal symbionts could elucidate whether evolutionary ancestors of the ELB took up fungal genes, which still benefit the descendants by e.g. encoding defensive compounds for the host beetles (Pankewitz and Hilker 2008), thus rendering them independent of harbouring resident, vertically transmitted fungi. Exploring such symbioses will improve our understanding of the evolution of these tripartite interactions between plants, phyllophagous beetles and microbes. This knowledge could further potentially lead to the development of more effective strategies for controlling ecologically and economically significant pests.

## Materials and Methods

### Insects and rearing conditions

Elm leaf beetles (ELB, *X. luteola*) were collected from a natural population in Montpellier, France, during summer 2021 and subsequently reared on potted, cloned elm trees (*U. minor,* three to four months old) in a greenhouse under long day conditions (18-hour light/6-hour dark cycle). Approximately 20 adult beetles were placed on leaves of an elm branch that was enclosed in a microperforated polypropylene bag, thus preventing escape of the beetles. Three times a week, branches were examined for egg depositions; then insects were transferred to fresh branches. Branches with egg clutches were also enclosed in bags. Hatchlings developed on the bagged tree branches until pupation. Pupae were transferred to aerated plastic containers in a climate-controlled chamber (18-hour light/6-hour dark cycle, 160 µmol m^-2^ s^-1^ light intensity, 20°C, 70% relative humidity) until adult emergence.

### Plant growth conditions

Elm trees (*U. minor*) were propagated using an *in vitro* shoot culture established from a single specimen from the Berlin Dahlem region, as described in Büchel *et al*. (2012). Once the trees had developed root systems, they were transferred to plastic pots containing a 3:1 soil-to-vermiculite mixture. These potted trees were kept in a climate-controlled chamber (22°C, 16-hour light/8-hour dark cycle, 160 µmol m^-2^ s^-1^ light intensity, 70% relative humidity). After ten weeks, the trees were transferred to a greenhouse, where they remained at long-day conditions until needed for experiments. Trees utilised in our study were approximately 14 to 15 weeks old.

### Microbial community sampling and analysis overview

To analyse the species richness and abundance of bacteria and fungi across ELB life stages and generations, freshly emerged adult beetle couples were placed on an elm branch of a tree for mating and egg deposition. The branch was then enclosed in a microperforated polypropylene bag to prevent the beetles from escaping (Figure 1). We subsequently sampled the parental (F_0_) couples and their laid eggs. For egg sampling, part of an egg clutch was gently removed from the lower surface of an elm leaf with sterilised tweezers. Larvae were allowed to hatch from the rest of the eggs. We collected samples from (F_1_) neonate larvae, seven-day-old larvae, pupae, and freshly emerged adults of both sexes. To obtain pupae and adults, prepupae were removed from the bags and transferred to sterile 2 mL reaction tubes with a pierced lid for further development at 25°C. The F_1_ pupae and resulting adult beetles had no contact to conspecifics or elms prior to sampling.

Each sample with F_0_ or F_1_ adult insects contained a single adult beetle, each sample with neonates contained a pool of 5 neonates, each sample with seven-day-old larvae contained a single individual larva, pupal samples contained each a single pupa, and egg samples contained 12-18 eggs.

To determine whether and how the bacterial and fungal communities of elm leaves match the microbial communities of *X. luteola,* we collected samples from intact elm leaves by cutting leaf sections with ethanol-sterilised metallic scissors. Each leaf sample consisted of 5 cm^2^ leaf material. Utilising ethanol-sterilised metallic forceps, we transferred the samples to 2 mL FastPrep^®^ tubes (Fisher Scientific). Moreover, we sampled feeding-damaged leaves to examine how feeding damage affects the microbial community associated with the leaves. These leaves were feeding-damaged by adult beetles for seven days. The size of these samples was equivalent to the size of samples from intact leaves.

We surface-sterilised parental F_0_ insects, F_1_ neonates, seven-day-old larvae, pupae, and freshly emerged F_1_ adults using sterilisation solution (0.5% v/v sodium hypochlorite, 0.1% v/v SDS, water). Eggs were not sterilised because microbes might be vertically transmitted from one generation to the next inside and outside the eggs. We added 500 µL sterilisation solution to the tubes containing the insect samples, vortexed the samples for 10 s, and rinsed them three times with autoclaved distilled water. This method effectively removes external microbes without affecting the internal microbial load, as demonstrated by a comparison of surface-sterilised insects with non-sterilised insects (data not shown). From this step on, sample processing was conducted in a biological safety cabinet to minimise contamination.

Sterile phosphate-buffered saline (PBS) was added to the tubes containing the samples. A volume of 150 µL was added to the egg and neonate samples, while 200 µL was added to all other samples. The samples were then bead-homogenised for 15 s at 4,500 rpm using a Precellys Evolution® tissue homogeniser.

For culture-dependent analyses of bacterial and fungal communities, 70 µL of the homogenate were processed immediately as described below. For culture-independent analysis of fungal communities, we stored the remaining volume at -80°C for subsequent further analysis.

For culture-independent analysis of bacterial communities, a separate set of samples was collected following the same experimental design (excluding feeding-damaged leaves) and stored at -80°C until further use. To account for potentially environmental contamination, negative control samples containing only PBS were processed in parallel with the experimental samples. Additionally, positive control samples were incorporated for the culture-independent analysis of bacterial communities. These positive controls consisted of PBS spiked with a resuspended pellet of *Escherichia coli* DH5-alpha (50 μL TE buffer), which had been pre-cultured in 1 mL of Lysogeny Broth (LB) at 37°C overnight.

### Culture-dependent analysis of microbial communities

To analyse bacterial and fungal communities via culture-dependent methods, homogenates were 1:10 serial diluted in PBS four times. Aliquots of 35 µL from each dilution were plated onto Tryptic Soy Agar (TSA) and Potato Dextrose Agar (PDA) supplemented with chloramphenicol (50 mg/L) (Figure 1). Colony-forming units (CFU) were counted after 48 hours at 27°C. Negative controls, consisting of PBS only, were performed to monitor for potential contamination.

CFU were morphologically characterised based on size, shape, colour, and texture, and subsequently restreaked to isolate pure cultures.

Genomic DNA extraction from pure cultures was performed by using the MasterPure™ DNA Purification Kit (Epicenter), following the manufacturer’s protocol. For samples designated for bacterial community analysis, an additional lysozyme digestion step was incorporated before the proteinase K digestion step to enhance bacterial lysis. This step involved the addition of 0.33 μL Ready-Lyse™ Lysozyme Solution, followed by a 15 min incubation period at room temperature.

Universal primers were used to amplify the 16S rRNA gene in bacterial isolates and the internal transcribed spacer (ITS) region in fungal isolates (Figure 1): 27F and 1492R for bacterial 16S rRNA gene (sequences: 5’-AGAGTTTGATCMTGGCTCAG-3’ and 5’-GGTTACCTTGTTACGACTT-3’, respectively), and ITS1 and ITS4 for fungal ITS region (sequences: 5’-TCCGTAGGTGAACCTGCGG-3’ and 5’-TCCTCCGCTTATTGATATGC-3’, respectively).

PCR analyses were carried out using the JumpStart Taq ReadyMix from Sigma-Aldrich, using 50 ng of DNA template in a reaction volume of 50 µL. Cycling parameters consisted of an initial denaturation cycle at 94°C for 2 min, followed by 30 cycles of denaturation at 94°C for 30 s, annealing at 52°C for 16S primers and 55°C for ITS primers for 30 s, extension at 72°C for 2 min, and a final extension cycle at 72°C for 5 min. The resulting PCR products were sent to Microsynth Seqlab, Germany, for Sanger sequencing.

For the microbial sequence identification, we used the Basic Local Alignment Search Tool (BLAST) accessible on the National Center for Biotechnology Information (NCBI) website (https://blast.ncbi.nlm.nih.gov/). The query sequences were compared against the NCBI database, and the top matches were analysed based on percent identity, alignment length, and E-value to identify the closest match to the query sequences. Isolates were given a strain name, and their sequences were deposited in the Sequenced Read Archive (SRA) database under the BioProject accession number PRJNA979994.

### Culture-independent analysis of microbial communities

For culture-independent analysis of the bacterial and fungal communities, genomic DNA was extracted from the above-described samples of elm leaves and different ELB life stages (Figure 1). The DNA extraction method was the same as described above tor the culture-dependent samples.

After DNA extraction, we amplified the bacterial 16S rRNA gene and fungal ITS region using universal primers optimised for the Illumina MiSeq platform. We employed 515F (5’-GTGYCAGCMGCCGCGGTAA-3’) and 806R (5’-GGACTACNVGGGTWTCTAAT-3’) primers for bacteria, as recommended by the Earth Microbiome Project (EMP: https://earthmicrobiome.org), and custom ITS primers developed by Usyk *et al*. (2017) for fungi: ITS1-30F (5’-GTCCCTGCCCTTTGTACACA-3’) and ITS1-217R (5’-TTTCGCTGCGTTCTTCATCG-3’). All primers included Illumina overhang adapter sequences for compatibility with index and sequencing adapters: forward adapter (5’-TCGTCGGCAGCGTCAGATGTGTATAAGAGACAG-3’) and reverse adapter (5’-GTCTCGTGGGCTCGGAGATGTGTATAAGAGACAG-3’).

For the PCR analyses, we used again the JumpStart Taq ReadyMix from Sigma Aldrich and 50 ng DNA template in a 50 µL reaction volume. PCR cycles were performed as described for the culture-dependent analysis. We visualised 10 µL of amplified product on a 1% agarose gel stained with ethidium bromide to assess target amplification. PCR amplicons were purified using MagBio HighPrep Clean-up magnetic beads (MagBio, USA), following the manufacturer’s protocol and added barcoded Illumina sequencing adapters. For this, a second PCR was performed using 5 μL purified PCR product, with initial denaturation at 95°C for 3 min, 8 cycles of 95°C for 30 s, 60°C for 30 s, 72°C for 30 s, and a final 72°C extension for 10 min. Indexed amplicons were purified using magnetic beads and quantified with a Qubit 2.0 fluorometer and the dsDNA high sensitivity assay kit (Thermo Scientific, USA).

Equimolar concentrations of each sample were pooled to create libraries. The final library’s quality and integrity were assessed using an Agilent 2200 TapeStation and D1000 ScreenTapes (Agilent Technologies, USA).

The combined library was sequenced at the Berlin Center for Genomics and Biodiversity Research (BeGenDiv) on the Illumina MiSeq platform, employing the MiSeq v3 (600 cycles) reagent kit for 2 × 300 bp paired-end reads.

### Sequence processing and analysis

The resulting data were analysed using a full-stack R pipeline (Callahan *et al*. 2016b) incorporating dada2 (Callahan *et al*. 2016a), phyloseq (McMurdie and Holmes 2013), and vegan (Oksanen *et al*. 2022). Adapter- and primer-trimmed reads were dereplicated and denoised using a parameterised model of substitution errors. The resulting denoised read pairs were merged and subjected to de novo chimera removal. Taxonomy was assigned using the latest Ribosomal Database Project training-set or UNITE for 16S and ITS, respectively. Bray-Curtis dissimilarity was calculated based on relative abundance to account for differences in library size and modelled using permutational multivariate ANOVAs.

### Insect performance assays

Spore suspensions of fungal isolates identified from ELB samples and elm leaves were prepared to determine how these fungi affect the insect’s performance parameters. Two predominant fungal isolates - designated as *Penicillium* sp. LPV01 and *Aspergillus* sp. LPV02 - were cultured on potato dextrose agar (PDA) with chloramphenicol (50 mg/L) at 27°C until sporulation (5-6 days). Spores were harvested by covering colonies with PBS and gently scraping the mycelium with a sterile inoculator. The spore-containing PBS was collected in sterile 50 mL Falcon tubes, filtered through sterile gauze, and centrifuged at 10,000 g for 2 min. The supernatant was discarded, and spores were resuspended in sterile water. The spore concentration was adjusted by transferring 500 µL of the suspension into an Eppendorf tube, vortexing, and pipetting a fixed volume into a Neubauer chamber for spore counting under a microscope. The spore suspension was diluted to 1000 spores/µL using sterile water. Aliquots of 50 mL were stored at 4°C until further use. Spore viability was regularly confirmed by plating spore samples on PDA before performing experiments.

To investigate the influence of *Penicillium* sp. LPV01 and *Aspergillus* sp. LPV02 on insect performance, we inoculated elm branches with either type of fungal spores. We collected 60-80 cm long branches from our greenhouse-grown trees, washed the leaves with sterile water, and surface-sterilised them by spraying them with 70% ethanol. Control (surface-sterilised) branches were left untreated, while branches for the *Penicillium* and *Aspergillus* treatment groups were sprayed with their respective spore suspensions (10^6^ spores/mL) until the entire surface was covered. The suspensions on the leaves dried at room temperature for three to five hours. Thereafter, the branches were individually placed in water-filled containers (25 mL), which were sealed with Parafilm, and then placed in plastic boxes labelled according to their treatment.

Neonate ELB larvae were randomly allocated to the treatment groups: (1) feeding on surface-sterilised leaves, (2) feeding on leaves inoculated with *Penicillium* spores, and (3) feeding on leaves inoculated with *Aspergillus* spores. Each treatment group consisted of 11-17 biological replicates, with each replicate containing five to eight neonates feeding together on the leaves of a branch. Larvae fed upon these leaves until pupation under standardised abiotic conditions (18-hour light/6-hour dark cycle, 160 µmol m^-2^ s^-1^ light intensity, 20°C, and 70% relative humidity). We recorded the survival rate of larvae after a feeding period of seven days and until pupation. Furthermore, we documented the pupal biomass using an analytical balance (Sartorius Lab Instruments GmbH & Co.). After emergence of the adult beetles, we paired individuals from the same treatment group and placed them on an untreated elm branch. For a period of two weeks, we then counted the number of eggs per egg clutch laid by females that had developed on the differently treated branches (Figure 2).

### Statistics

Statistical analysis was conducted in R (version 4.2.1) for bacterial quantification and insect performance data. Normality of the data was evaluated by Shapiro-Wilk test, and variance homogeneity was checked by Levene’s test. Parametric and non-parametric tests were chosen based on the distribution of the data. We used the Kruskal-Wallis (KW) test followed by Dunn’s multiple-comparison test with Benjamini-Hochberg *post hoc* (Dunn-BH) correction for multiple comparisons. Pairwise comparisons were analysed using Student’s *t*-test. Principal-coordinate (PCo) analysis on Bray-Curtis dissimilarity and PERMANOVA was to analyse microbial beta diversity.

## Data availability

The sequences corresponding to the 16S rRNA gene and the ITS region from the bacterial and fungal isolates are available in GenBank under the accession numbers OR136186-OR136188 and OR136192-OR136207. The data obtained from Illumina sequencing has been uploaded to the SRA database with the Bioproject accession number PRJNA979994.

## Supplemental material

Supplemental material is available online.

FIGURE S1. Overview of bacteria detection via PCR

TABLE S1: Analysis of bacterial isolates via BLAST

TABLE S2. Analysis of fungal isolates via BLAST

## Acknowledgements

We thank Gabriele Haberberger and Laura Hagemann at the Freie Universität Berlin for their assistance in growing plants and rearing the insects. This study was supported by the German Research Foundation (Deutsche Forschungsgemeinschaft) (Collaborative Research Centre 973, project B1; http://www.sfb973.de/).

J.S., M.H., M.R.E. and L.R.P.V. conceptualised the study. J.S., J.R. and L.R.P.V performed the experiments. L.R.P.V., S.M. and S.P. processed the samples for MiSeq sequencing. P.J. performed the sequencing analysis. L.R.P.V evaluated the ecological and microbiological data. J.S. and L.R.P.V. wrote a first draft of the manuscript. All authors contributed to later versions of the manuscript and agreed with the final version.

